# APOBEC3A-Induced DNA Damage Drives Polymerase θ Dependency and Synthetic Lethality in Cancer

**DOI:** 10.1101/2025.10.21.682588

**Authors:** Abhishek Bose, Weisi Liu, Paul Yoo, Alina Sami, Michal T. Boniecki, Madhura Deshpande, Mohamed Osman, Duy Nguyen, Urko D. Castillo, Paraskevi Giannakakou, Bhavneet Bhinder, William Hooper, Olivier Elemento, Nicolas Robine, Milaid Granadillo Rodriguez, Jeannine Gerhardt, Kent W. Mouw, Linda Chelico, Bishoy M. Faltas

**Author notes:** **Corresponding Author:** Bishoy M. Faltas. These authors equally contributed to this manuscript (Abhishek Bose, Weisi Liu). These authors co-supervised the work (Kent W. Mouw, Linda Chelico, Bishoy M. Faltas).

## Abstract

APOBEC3 cytidine deaminases drive cancer evolution. There is an unmet need to target cancer cells with APOBEC3 activity. Here, we identify error-prone theta-mediated end joining (TMEJ) as the main pathway for repairing APOBEC3-induced double-strand breaks (DSBs). Using fluorescent DSB repair reporters and a novel biochemical assay, we demonstrate that APOBEC3A competes with replication protein A (RPA) for single-stranded DNA overhangs, exposing microhomologous sequences to shift DSB repair towards error-prone TMEJ. Genomic analysis of clinical tumor samples confirmed the co-occurrence and proximity between APOBEC3-induced mutational footprints, microhomology-mediated deletions (MMDs), and TMEJ-associated chromosomal instability signatures. Crucially, inhibition of DNA polymerase theta (Polθ) synergizes with APOBEC3A-induced DSBs to induce synthetic lethality *in vitro* and *in vivo*. Collectively, our findings identify TMEJ as the preferred mechanism for repairing APOBEC3A-induced DSBs and establish Polθ inhibition as a novel promising strategy to eliminate cancer cells with APOBEC3A activity.

## Introduction

The APOBEC3 family of cytosine deaminases plays a pivotal role in eukaryotic immune defense by restricting viral replication (1,2). However, tumor cells hijack these endogenous mutators, particularly APOBEC3A and APOBEC3B, to fuel genetic diversification and to adapt rapidly under selective pressures (3-5). APOBEC3-driven mutagenesis is prevalent across various cancers (6-8), conferring critical evolutionary advantages, including clonal mutagenesis, and resistance to several anticancer therapies (9-13). Recent studies increasingly implicate APOBEC3A as a key driver of mutagenesis and therapy resistance in several cancers, including urothelial carcinoma (14-19). Although efforts to inhibit APOBEC3 enzymes are underway, no clinical-stage inhibitors have been reported to date (15,20-22).

Prior studies have shown that APOBEC3A-induced lesions in single-stranded DNA (ssDNA) are protected by mechanisms that safeguard replication-fork integrity; however, APOBEC3-induced cytosine deamination ultimately leads to double-strand breaks (DSBs) and chromosomal instability (23-26). In this study, we investigate how cancer cells withstand potentially lethal damage from APOBEC3A-induced DSBs. We discovered that TMEJ, a double-strand break (DSB) repair pathway dependent on Polθ and short microhomology sequences at break ends, is preferentially activated under APOBEC3A activity conditions. We show that APOBEC3A-induced DSBs are repaired by error-prone TMEJ, contributing to chromosomal instability in patients with urothelial carcinoma. Crucially, we demonstrate that inhibiting Polθ in urothelial carcinoma cells with APOBEC3A activity induces synthetic lethality *in vitro* and *in vivo*. Together, this study presents a promising therapeutic avenue to exploit vulnerabilities arising from APOBEC3A-induced DSBs to target cancer cells to counter tumor evolution and therapy resistance in the clinic.

## Results

### APOBEC3A induces replication-fork stalling and double-strand DNA breaks

APOBEC3A-induced mutagenesis occurs in episodic bursts that rapidly elevate mutation rates over short intervals (27,28), providing tumor cells with evolutionary advantages yet causing severe genomic damage. To recreate this dynamic pattern, we employed a doxycycline-inducible APOBEC3A system in the 5637 bladder cancer cell line, a well-characterized basal-type bladder cancer cell line (29,30). We hypothesized that upon ssDNA binding, APOBEC3A-induced deamination induces replication-fork stalling, thereby activating CHK1 signaling. Indeed, following doxycycline induction, we observed that APOBEC3A expression led to increased phosphorylation of CHK1 (Ser345) and replication protein A (RPA) (Ser4/8), as well as elevated γH2AX and CHK2 (Thr68), indicating replication stress and DSB formation over time (**Fig. 1A**). APOBEC3A expression also increased cleaved PARP1 (Asp214), a marker of cellular apoptosis (31) (**Fig. 1A**). These results suggest that APOBEC3A bursts trigger a DNA damage cascade involving replication-forks stalling, DSB formation, and eventually apoptosis due to accumulated unrepaired damage.

**Figure 1.**
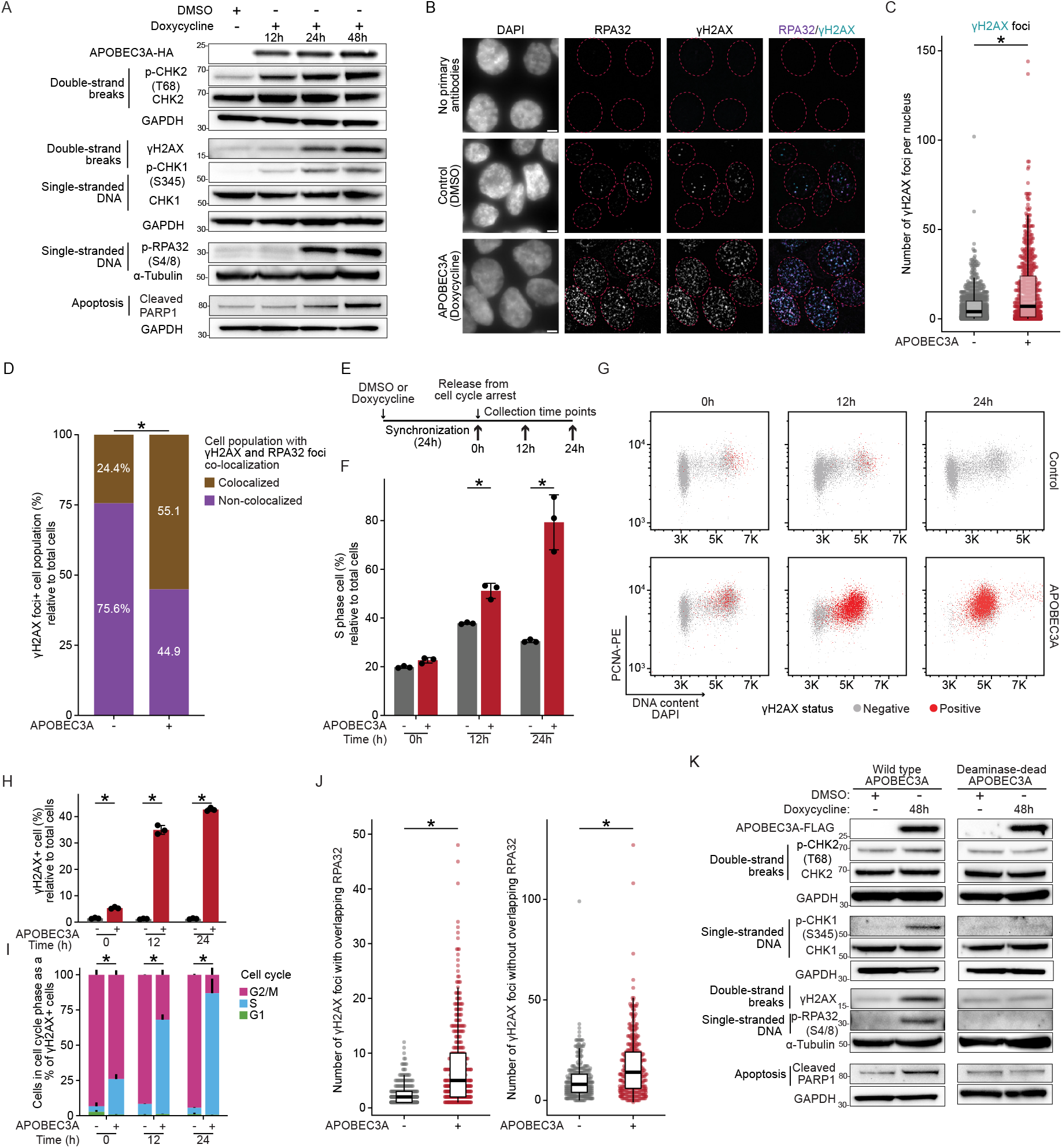
APOBEC3A-induced mutation bursts cause double-strand breaks, replication stress, and cell cycle arrest in S phase. A, SDS-PAGE western blot showing that induction of APOBEC3A-HA in 5637 urothelial carcinoma cells, initiates a DNA damage cascade, with concurrent accumulation of ssDNA markers p-RPA(Ser4/8) and p-CHK1 (Ser345); DSBs markers γH2AX and p-CHK2 (Thr68); and resulting cellular apoptosis indicated by increased cleaved PARP1 from 12 to 48 hours. Blots with their individual loading controls are separated by a gap. B, Representative immunofluorescence images of 5637 cells treated with doxycycline (APOBEC3A induction) or DMSO (control) for 24 hours, stained for RPA32 and γH2AX foci in the nuclei. Scale bar = 5 μm. Dotted line marks the nuclear boundary. C, Quantitative box plot with individual data points illustrating the significant increases in the number of γH2AX foci per nucleus upon APOBEC3A induction. Each dot represents one nucleus. Two-sided t-test. D, Bar chart illustrating the percentage of γH2AX foci-positive nuclei that have overlapped RPA32 and γH2AX foci or γH2AX foci alone, indicating that cells with colocalized RPA32 and γH2AX foci are enriched in the APOBEC3A-expressing population compared to controls. Fisher’s exact test. E, Schematic of the experimental design for the cell cycle and exposure to APOBEC3A. F, Quantitative bar chart illustrating that APOBEC3A causes cell-cycle arrest in S phase, multiple t-test. G, Scatter plot for the cells with cell cycle (DAPI and PCNA) and γH2AX staining, showing cell cycle distribution and concurrent DNA damage. H, Bar chart illustrating the percentage of γH2AX-positive cells relative to total cells, indicating APOBEC3A significantly increased the γH2AX-positive cells. Multiple t-test. I, Bar chart illustrating the distribution of γH2AX-positive cells across different cell cycle stages with a significant enrichment in S-phase cells, Fisher’s exact test. J, Scatter plot with boxplot illustrating the number of RPA and γH2AX co-localization foci (ends with ssDNA overhangs) and γH2AX foci without RPA32 foci (newly formed two-ended breaks) in the S-phase arrested cells (nuclei positive for co-localization of RPA32 and γH2AX), indicating that APOBEC3A significantly increased both break events compared to DMSO controls. K, SDS-PAGE western blot analysis comparing the DNA damage cascade induced by wildtype APOBEC3A and the deaminase-inactive APOBEC3A E72Q variant in RT112/84 urothelial carcinoma cells after 48 hours of doxycycline induction. Wildtype APOBEC3A significantly induced accumulation of ssDNA markers p-RPA(Ser4/8) and p-CHK1 (Ser345); DSBs markers γH2AX and p-CHK2 (Thr68); and resulting cellular apoptosis indicated by cleaved PARP1, while the deaminase-inactive variant did not elicit these effects. Blots with their individual loading controls are separated by a gap. *: p < 0.05. Box plots show the median and IQR; the lower whisker indicates Q1–1.5 × the IQR; the upper whisker indicates Q3 + 1.5× the IQR.

During DNA replication, exposed ssDNA is highly susceptible to APOBEC3A-induced deamination (24). The resulting uracil or the subsequent abasic site generated by uracil N-glycosylase (UNG) removal of uracil can impede replicative polymerase progression, leading to replication-fork stalling (**Supplementary Fig. S1**) (32). To test this hypothesis, we performed DNA fiber assays that track replication-fork dynamics (33). Cells were pre-treated with DMSO or doxycycline to induce APOBEC3A expression for 24 hours in serum-free medium, during which APOBEC3A-induced DNA damage accumulated while cells were largely outside S phase. Upon release into the S phase by serum reintroduction, replication forks encountered this pre-existing damage. After releasing cells into S phase by reintroducing serum, we sequentially labeled replicating DNA with two nucleotide analogs, CldU followed by IdU, in the presence or absence of the DNA replication inhibitor aphidicolin. This sequential labelling enabled monitoring of replication-fork progression by comparing CldU and IdU tract lengths, with the CldU pulse reflecting baseline progression under controlled APOBEC3A exposure. Because APOBEC3A expression persisted during both labeling pulses, additional lesions accumulated as labeling progressed. APOBEC3A expression significantly decreased the ratio of nascent DNA tract lengths (median IdU/CldU ratio: 4.6 in APOBEC3A-expressing cells vs. 7.2 in DMSO controls, p < 0.0001). As expected, DNA polymerase inhibitor aphidicolin-treated cells showed a significant reduction (median: 4.6, p < 0.0001) in track ratio compared with DMSO-treated controls. Importantly, aphidicolin-treated cells exposed to APOBEC3A exhibited the greatest reduction in tract ratio among all, (median: 3.3, p < 0.0001) with a significant difference compared to aphidicolin-treated cells alone (**Supplementary Fig. S2A and S2B**). This additive effect reflects how APOBEC3A exacerbates replication-fork slowing caused by low-dose aphidicolin treatment. Collectively, these findings confirm that APOBEC3A induces replication-fork stalling consistent with the observed CHK1 activation. (**Fig. 1A)**.

APOBEC3A preferentially deaminates cytosines on the lagging strand during DNA replication (34), potentially resulting in two-ended DSBs through UNG activity and subsequent apurinic/apyrimidinic endonuclease-mediated excision (**Supplementary Fig. S1)** (35-37). To characterize the overlap of DNA break ends with ssDNA overhangs within individual nuclei, we used immunofluorescence microscopy. We quantified the colocalization of RPA (38) and γH2AX foci (39) (**Fig. 1B**). After 24 hours of APOBEC3A induction, the total number of γH2AX foci per nucleus increased significantly (mean: 15.2 in APOBEC3A-expressing cells vs. 6.5 in DMSO controls, p < 0.001) (**Fig. 1C**), indicating extensive DSB formation. Moreover, the percentage of cells with colocalization of γH2AX and RPA, increased in APOBEC3A-expressing cells compared to controls (Fisher’s exact test, p < 0.001) (**Fig. 1D**). Since ssDNA overhangs typically arise via end-resection or replication-fork collapse in the S phase (40,41), we hypothesized that APOBEC3A-induced DSBs arrest the cells in S phase due to the accumulation of resected breaks with ssDNA overhangs that serve as substrates for subsequent DSB repair.

We quantified the S-phase marker PCNA, alongside γH2AX staining by flow cytometry (**Fig. 1E**). After 24 hours of cell cycle release post-synchronization, APOBEC3A-expressing cells displayed a pronounced S-phase arrest compared to DMSO controls (p < 0.001) (**Fig. 1F and Supplementary Fig. S2C**). This S-phase arrest was associated with a marked accumulation of APOBEC3A-induced DSBs (mean: 42% of γH2AX-positive cells in APOBEC3A-expressing cells vs. 1.2% in DMSO controls, p < 0.001) (**Fig. 1G and 1H**). Also, the percentage of cells in S phase was significantly higher in APOBEC3A-induced γH2AX-positive cells compared to the DMSO treated cells (Fisher’s exact test, p < 0.001) (**Fig. 1I**). APOBEC3A significantly increased nuclei with co-localized γH2AX and RPA foci indicative of DSBs with ssDNA overhangs (mean: 17 in APOBEC3A-expressing cells vs. 9.9 in DMSO controls, p < 0.001). Furthermore, nuclei with DSB (γH2AX) not co-localized with RPA, indicative of unresected DSBs without ssDNA overhangs, were also higher in APOBEC3A-expressing cells (mean: 7.5) compared to 2.3 in controls (p < 0.001), suggesting an increase in DSBs without ssDNA overhangs (**Fig. 1J**). These findings indicate that APOBEC3A activity substantially enhances the generation of DSB with and without ssDNA overhangs, resulting in S-phase arrest. To confirm that APOBEC3A’s DNA-damaging effects depend on its deaminase activity across different bladder cancer subtypes, we compared wild-type and deaminase-inactive APOBEC3A variants in the RT112/84 cells, a luminal cell line with low baseline genome instability (29,30). Only wild-type APOBEC3A activated CHK1/CHK2, increased RPA phosphorylation and γH2AX levels, and induced PARP cleavage (**Fig. 1K**). These results demonstrate that the APOBEC3A-induced DNA damage cascade depends on its deaminase activity and occurs across molecularly distinct bladder cancer backgrounds.

### Microhomology-mediated deletions and chromosomal instability signatures correlate with APOBEC3 activity in human cancers

Cancer cells employ multiple repair mechanisms to repair potentially catastrophic DSBs, including homologous recombination (HR), classical non-homologous end joining (c-NHEJ), and theta-mediated end joining (TMEJ) (40). TMEJ serves as a backup mechanism to HR during the S-phase, using microhomology sequences within ssDNA overhangs generated during the end resection step of HR (40,41). TMEJ is also the central repair mechanism for persistent DSBs that originate in the S phase and persist into mitosis (42,43). Conversely, c-NHEJ activity is more active during the G1 phase, as it primarily repairs blunt or minimally processed DNA ends. In contrast, its activity is reduced in S and G2 phases due to increased end resection and targeted degradation of the Ku70-Ku80 complex by E3 ubiquitin ligases (40,44). Since APOBEC3A activity-induced DSB formation causes S-phase arrest **(Fig. 1E)** (24), we hypothesized that either HR or TMEJ serves as the primary repair mechanism for APOBEC3A-induced DSBs.

To investigate whether TMEJ repairs APOBEC3A-induced DSBs, we quantified microhomology-mediated deletions (MMDs) following inducible APOBEC3A expression **(Fig. 2)**. These deletions potentially arise from inappropriate end-joining (45) caused by uneven annealing of microhomology sequences during TMEJ, leading to excision of intervening DNA (8,46). We analyzed whole-exome sequencing data from an isogenic 5637 cells exposed intermittently to 6 cycles of doxycycline-regulated APOBEC3A recapitulating previously described episodic expression patterns (28) compared to non-exposed controls (**Methods**) (**Fig. 2A**). We used the Catalogue of Somatic Mutations in Cancer (COSMIC) (47) framework to detect deletion signatures adjacent to microhomology sequences at break ends based on the COSMIC annotations of signatures representing TMEJ activity (8,46). We found that intermittent exposure to APOBEC3A significantly elevated the burden of MMDs, with a mean of 13 MMDs in APOBEC3A-expressing cells (replicate n=5) compared to 9 in DMSO controls (replicate n=3) (p = 0.04) (**Fig. 2B; Supplementary Table S1**). These data confirm that TMEJ-mediated MMDs accumulate following episodic APOBEC3A expression.

**Figure 2.**
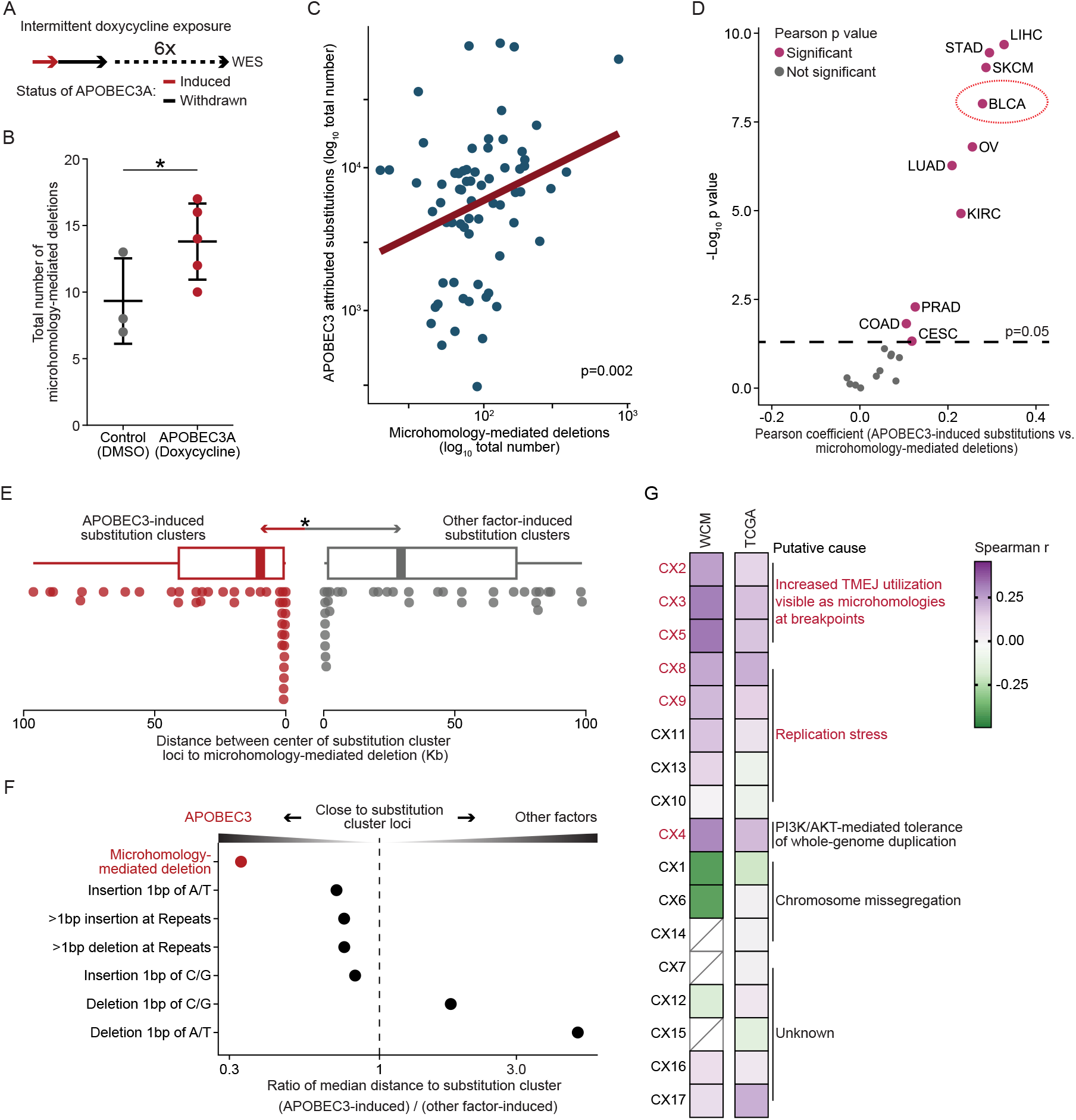
APOBEC3-induced mutagenesis is associated with microhomology-mediated deletions and chromosomal instability. A, Schematic of the experimental design for long-term intermittent exposure of 5637 cells to APOBEC3A. B, Dot plot illustrating that intermittent APOBEC3A exposure significantly increased the number of MMDs. Each dot represented one replicate. One-sided t-test. Data represent mean ± SD. C, Scatter plot showing a significant positive correlation between the burdens of APOBEC3-induced single-base substitutions and MMDs in whole-genome sequencing data from 67 urothelial carcinoma samples. Pearson r=0.36. D, Dot plot illustrating the Pearson correlation coefficients and corresponding p-value across TCGA pan-cancer cohorts, indicating consistent associations between the burdens of APOBEC3-induced substitutions and MMDs across various cancers. E, Box plot showing the distances of MMDs to the center of substitution clusters. Each dot indicates the distance between MMDs and its corresponding substitution cluster. One-sided t-test. Box plots show the median and IQR; the lower whisker indicates Q1–1.5 × the IQR; the upper whisker indicates Q3 + 1.5× the IQR. F, Dot plot illustrating the ratio of the median distance of indel footprints to APOBEC3-induced substitution clusters normalized to substitution clusters induced by other factors. MMDs were found to be closer to APOBEC3-induced substitution clusters, suggesting that TMEJ is preferentially engaged in repairing DSBs within regions affected by APOBEC3 activity. G, Heatmap illustrating the correlations between the burdens of APOBEC3-induced substitutions and chromosomal instability signatures, indicating that APOBEC3-induced substitutions are significantly correlated to signatures related to replication stress and increased TMEJ repair. Red fonts indicate Spearman p < 0.05 in either the WCM or BLCA cohort. CIN values below detection thresholds are indicated as boxes with diagonal lines.

To evaluate whether human cancer genomes recapitulate these *in vitro* observations, we analyzed MMDs in whole-genome sequencing data from our previously published Weill Cornell Medicine (WCM) cohort of urothelial carcinomas (n=67) (**Methods**)(19). We identified a significant positive correlation between APOBEC3-induced substitutions (C>T/G in the TCW motif, where W=A/T) and MMDs in these patient samples (Pearson r=0.36, p = 0.002) (**Fig. 2C; Supplementary Table S2**). We further confirmed these observations in whole-exome sequencing data from The Cancer Genome Atlas (TCGA) pan-cancer cohort (n=8836). APOBEC3-induced substitutions showed a consistent correlation between MMDs across multiple cancer types, including hepatocellular carcinoma (Pearson r = 0.32, p = 2.09 x 10^-10^), stomach adenocarcinoma (Pearson r = 0.29, p = 3.53 x 10^-10^), and urothelial carcinoma (Pearson r = 0.27, p = 9.7 x 10^-09^) (**Fig. 2D)**. Previous studies have identified that mutational clusters caused by APOBEC3-induced processive activity known as kataegis frequently occurring near DSB sites (48-50). In our WCM patient cohort, we discovered that MMDs were significantly closer to APOBEC3-induced substitution clusters (enriched with coordinated C>T/G in the TCW motif, where W=A/T) (mean distance: 0.024Mb) compared to clusters caused by other mutagenic factors, such as chemotherapy (mean distance: 0.037Mb, p = 0.04). Other indel types not related to TMEJ did not exhibit similar proximity to kataegis clusters **(Fig. 2E, 2F; Supplementary Table S3)**. Given the coupling and spatial proximity between APOBEC3-induced substitutions and MMDs, we hypothesized that APOBEC3A not only induces DSBs but also biases repair pathway selection toward TMEJ. DSBs threaten genomic integrity and can cause chromosomal instability through large- and small-scale deletions, loss of heterozygosity, and translocations (51). We examined whether APOBEC3 activity is associated with complex chromosomal instability resulting from the interplay between APOBEC3-induced DSBs and TMEJ repair. We applied a computational framework (52) that characterizes 17 chromosomal instability signatures to the Weill Cornell Medicine (WCM) urothelial tumors (n=67) and TCGA Bladder Urothelial Carcinoma (BLCA) dataset (n=247) (**Methods**). In both cohorts, APOBEC3-induced C to T/G in the TCW motif (W=A/T) showed a significantly positive correlation with signatures associated with replication stress (CX8 and CX9) (p < 0.05), and impaired homologous recombination (IHR), namely CX2 (p < 0.05), CX3 (p < 0.01), and CX5 (p < 0.05). CX2 and CX3 are associated with increased utilization of the TMEJ and single-strand annealing (SSA) repair pathways, as evident from the presence of microhomologies at the breakpoints (52) (**Fig. 2G; Supplementary Table S4**). Collectively, these analyses further support for the critical role for TMEJ in repairing DSBs within regions affected by APOBEC3-induced mutagenesis, leaving detectable MMDs and specific chromosomal instability signatures as residual footprints from inappropriate end-joining.

### APOBEC3A binding to ssDNA overhangs displaces RPA and shifts DSB repair choice towards error-prone TMEJ

Given the coupling and spatial proximity between APOBEC3-induced substitutions and MMDs, we hypothesized that APOBEC3A induces DSBs and biases repair pathway selection toward TMEJ. We tested this using DSB repair reporters in U2OS cells, quantifying HR and TMEJ outcomes by measuring GFP restoration following I-SceI-induced DSB repair (**Fig. 3A; Supplementary Fig. S3A**). The DR-GFP reporter measures HR efficiency using a homologous template to repair DSBs and restore GFP expression accurately (53). The EJ2-GFP reporter, specifically designed for quantifying TMEJ efficiency, requires annealing of microhomology sequences near the break site to restore GFP expression (54) (**Methods**). Notably, APOBEC3A induction significantly decreased HR (p = 0.03) (**Fig. 3B**), while enhancing TMEJ (p = 0.006) (**Fig. 3C**). These results suggest that APOBEC3A shifts repair towards TMEJ for repairing DSBs.

**Figure 3.**
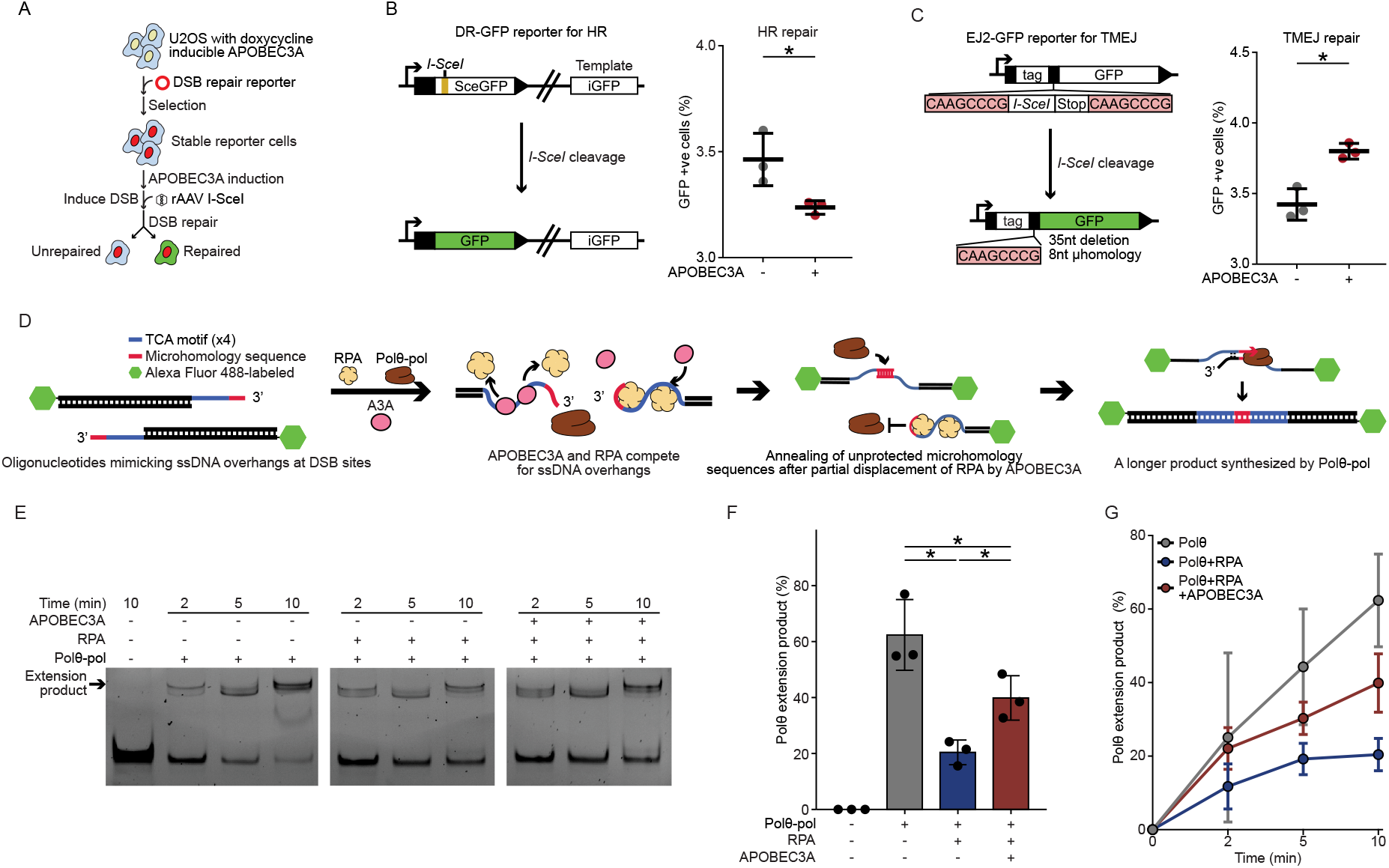
APOBEC3A competes with RPA for ssDNA overhangs, promoting TMEJ repair. A, Schematic workflow for the DSB reporter assay experiment used to assess the impact of APOBEC3A on DNA repair pathways. B, Schematic representation of the DR-GFP reporter assay for HR (Left). The accompanying dot plot (Right) illustrates the percentage of GFP-positive cells, indicating restoration of GFP via HR. The data show that APOBEC3A significantly reduced the HR efficiency in U2OS cells. Two-sided t-test. Data represent mean ± SD. C, Schematic representation of the EJ2 reporter assay specific for TMEJ (Left). The accompanying bar chart (Right) illustrates the percentage of GFP-positive cells, indicating restoration of GFP via TMEJ. The data show that APOBEC3A significantly increased the TMEJ efficiency in U2OS cells. Two-sided t-test. Data represent mean ± SD. D, Schematic of the competition assay for Polθ-mediated extension (TMEJ assay). Fluorescently labeled complementary oligonucleotides were used to simulate DSB ends with ssDNA overhangs, incorporating APOBEC3A-preferred motifs and microhomology sequence for annealing. Purified RPA, APOBEC3A, and the polymerase domain of Polθ (Polθ-pol) were introduced to check competition between APOBEC3A and RPA for the ssDNA overhangs, and its effect on the extension product synthesized by Polθ was quantified from gel electrophoresis of the product. E, Representative gel images illustrating the time-dependent extension products synthesized by Polθ-pol alone, Polθ-pol with RPA, and Polθ-pol with both RPA and APOBEC3A. F, Quantitative bar chart illustrating the Polθ efficiency of extension product under each condition at 10 minutes, indicating RPA binding significantly decreased Polθ-mediated synthesis, while the presence of APOBEC3A counteracted RPA and significantly enhanced Polθ-mediated synthesis. Each dot represents one independent experiment. Two-sided t-test. Data represent mean ± SD. G, Quantitative dot plot with error bars illustrating the time-dependent efficiency of Polθ-pol mediated extension product under each condition. Data represent mean ± SD. Three independent experimental replicates were performed for each condition. *: p < 0.05.

In addition to targeting ssDNA during replication, APOBEC3A also binds to and deaminates ssDNA overhangs formed at DSB ends (48-50). We hypothesized that APOBEC3A-RPA competition for ssDNA overhangs generated by end resection at DSBs is the mechanism by which APOBEC3A shifts DSB repair pathway choice towards TMEJ. This competition reduces RPA loading on ssDNA, thus exposing microhomology sequences to enable strand annealing and subsequent Polθ-dependent fill-in synthesis (55,56). To test this hypothesis, we modified an existing biochemical assay to recreate the competitive binding between APOBEC3A and RPA on ssDNA overhangs, allowing us to evaluate how this competition affects Polθ-mediated oligonucleotide extension (57) (**Methods**) (**Fig. 3D)**. We used fluorescently labeled complementary oligonucleotides that mimic ssDNA overhangs at DSB ends and incorporating APOBEC3A-preferred trinucleotide motifs (TCA) with adjacent microhomology sequences to enable annealing. Upon introducing purified RPA, the polymerase domain of Polθ (Polθ-pol) (58), and APOBEC3A proteins at a 1:1:2 ratio to these ssDNA oligonucleotide substrates, we found that the presence of RPA significantly reduced Polθ-mediated synthesis efficiency from 62% (Polθ-pol alone) to 20% (p = 0.005), calculated as percentage of the total band intensity per lane (substrate and product combined). Notably, the addition of APOBEC3A in the presence of RPA significantly rescued Polθ-mediated synthesis efficiency, restoring it to 39% (p = 0.02) (**Fig. 3E, 3F**). These data suggest that APOBEC3A facilitates TMEJ by competing with RPA for binding to ssDNA overhangs, thereby reducing RPA occupancy and exposing microhomology sequences for annealing and Polθ-mediated extension. Moreover, the increasing TMEJ efficiency was time-dependent (**Fig. 3G)**, indicating that the APOBEC3A-RPA competition was not instantaneous, but a dynamic process governed by the changing balance of APOBEC3A and RPA binding to ssDNA overhangs. These findings define a new mechanism by which APOBEC3A promotes TMEJ over alternative DSB repair pathways through competition with RPA for ssDNA overhangs.

### Targeting Polθ induces synthetic lethality in cancer cells expressing APOBEC3A

Building on our discovery that cancer cells depend on TMEJ to repair APOBEC3-induced double-strand breaks, we developed a therapeutic strategy targeting this dependency. We hypothesized that Polθ inhibition in APOBEC3-high cancer cells would disrupt their primary DNA repair mechanism, amplify genomic instability, and trigger cell death through accumulated DNA damage. To test this hypothesis, we used shRNA to knock down Polθ in doxycycline-inducible APOBEC3A-expressing 5637 cells (**Supplementary Fig. S3B**). Upon APOBEC3A induction, Polθ knockdown cells exhibited a significant 20% reduction in viable cell number compared to scramble shRNA controls (p < 0.001) (**Fig. 4A**) and increased apoptosis (**Fig. 4B)**. To confirm this relationship with endogenous APOBEC3A expression, we analyzed Chronos dependency scores from DepMap project, which measures gene fitness effects derived from CRISPR/Cas9 knockout screens across a panel of urothelial cancer cell lines (**Methods**) (59). We found a significant correlation between APOBEC3-induced substitutions and Polθ dependency scores in urothelial carcinoma cell lines (Spearman r = 0.41, p = 0.01, n=25) (**Fig. 4C; Supplementary Table S5**). These data confirmed that elevated APOBEC3 activity is associated with increased reliance on Polθ for survival.

**Figure 4.**
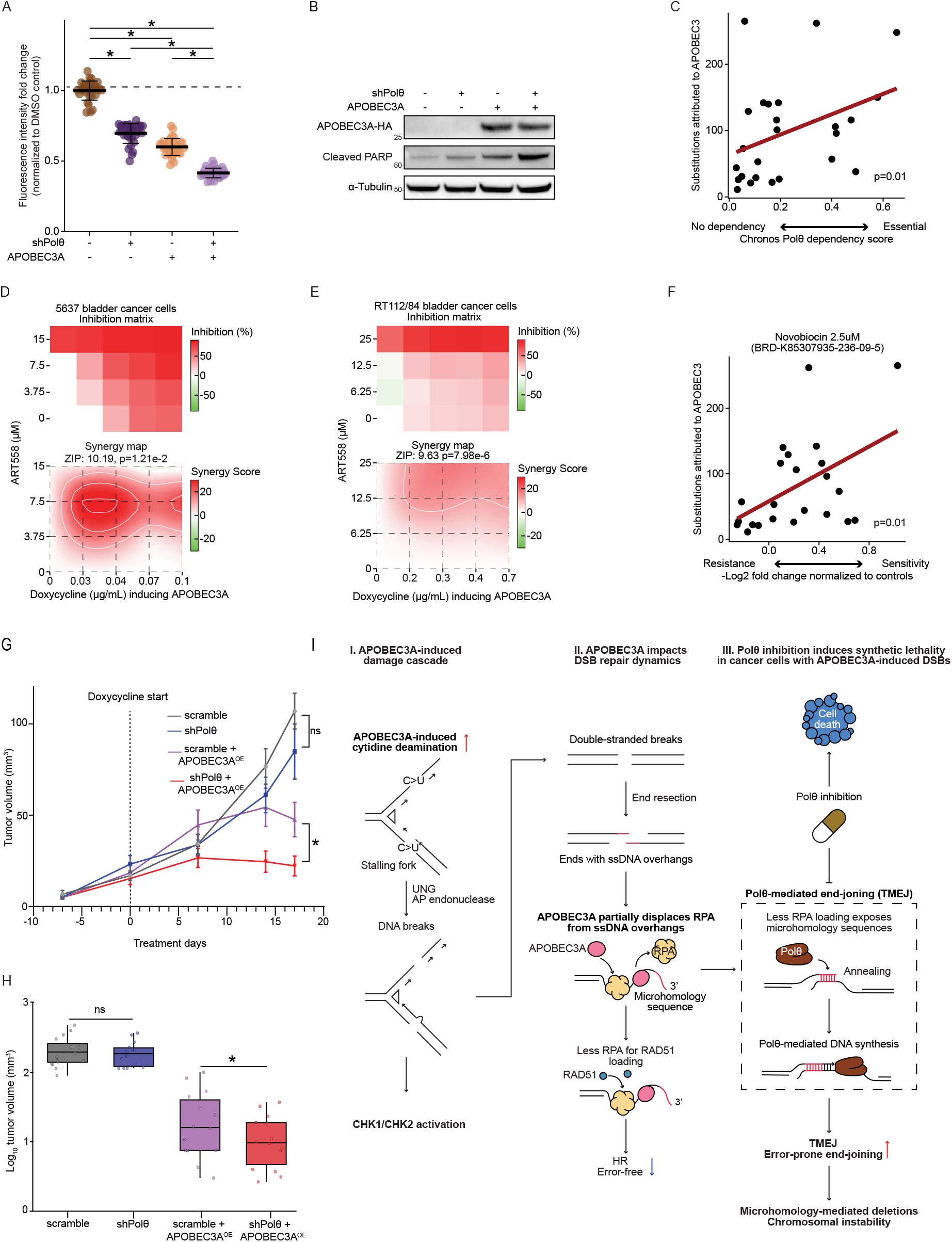
Targeting Polθ induces synthetic lethality with APOBEC3 activity *in vitro* and *in vivo*. A, Dot plot illustrating significantly decreased cell proliferation in Polθ knockdown cells compared to scramble controls with or without APOBEC3A induction after 48 hours. Each dot represented one replicate. Multiple two-sided t-tests. Data represent mean ± SD. B, SDS-PAGE western blot showing that APOBEC3A induction leads to significantly more cellular apoptosis in Polθ knockdown cells compared to scramble controls, indicated by cleaved PARP1. C, Scatter plot showing a significant correlation between APOBEC3-induced substitutions and the genetic dependency of urothelial carcinoma cells on Polθ, indicating that cells with higher APOBEC3 activity rely more on Polθ function for survival (n=25, Spearman r = 0.41). D and E, Heatmaps illustrating the inhibition matrix and synergistic interaction between APOBEC3A expression and the Polθ inhibitor ART558 in 5637 (D) and RT112/84 (E) urothelial cancer cells. Percent inhibition represents the reduction in cell proliferation in doxycycline-treated cells compared to DMSO controls. Dark red regions indicate stronger inhibition, while lower values are shown in lighter shades. For synergistic interaction, a heatmap showing red areas indicate synergistic effects, and green areas indicate antagonistic effects. The synergistic score was calculated using the ZIP model, and corresponding p-values for the significance of the synergistic effect was obtained using SynergyFinder Plus. F, Scatter plot showing a significant correlation between APOBEC3-induced substitutions and sensitivity to Novobiocin, a first-in-class Polθ inhibitor, in urothelial cancer cell lines. This indicates that pharmacological inhibition of Polθ increases the cytotoxicity induced by endogenous APOBEC3 activity. (n=23, Spearman r = 0.46, *: p < 0.05). G, Tumor growth kinetics showing mean tumor volume (± SEM) measured *in vivo*. NSG mice were injected first bilaterally with 5637 bladder cancer xenografts stably expressing scrambled or shPolθ shRNA and subsequently, at day 35 post-implantation (Day 0 of treatment), treated with either PBS (vehicle) or doxycycline (50mg/kg body weight) for APOBEC3A overexpression (APOBEC3A^OE^). Treatment was continued on alternate days in a weekly regimen (days 1, 3, 5, and 8) for 2 weeks. At each measurement time point (Day 7, 14) and at endpoint (Day 17 post-treatment), the shPolθ +APOBEC3A^OE^ group exhibited a significant tumor growth suppression relative to the scramble+APOBEC3A^OE^. PBS vehicle-treated groups (scramble vs shPolθ) had no significant (ns) difference (two-sided Student’s t-test). *: p < 0.05. H, Box plots showing endpoint log_10_-transformed tumor volumes ex vivo. shPolθ +APOBEC3A^OE^ group showed a significant reduction in tumor volume compared to scramble+APOBEC3A^OE^. PBS-treated vehicle groups showed a non-significant (ns) difference (one-sided Student’s t-test). Box plots represent median (center line), interquartile range (box limits), and 1.5× IQR (whiskers). n=16 tumors (8 mice × 2 tumors): scrambled (PBS) and scramble+APOBEC3A^OE^ (Doxycycline); n=14 tumors (7 mice × 2 tumors): shPolθ (PBS) and shPolθ +APOBEC3A^OE^ (Doxycycline). Each dot represents one tumor. Tumor volumes are plotted on a log10-transformed scale for visual clarity. Statistical significance was assessed using raw, untransformed values. *: p < 0.05. I, Schematic mechanistic model illustrating how APOBEC3A-induced mutagenic bursts generate extensive ssDNA during replication, leading to replication-fork stalling and DSB formation (activating CHK1 and CHK2). APOBEC3A then competes with RPA for ssDNA binding, limiting RAD51-mediated homologous recombination. Unprotected ssDNA exposes microhomology sequences, enabling error-prone TMEJ by Polθ. Inhibition of Polθ further exacerbates DNA damage, causing synthetic lethality in cells with elevated APOBEC3A, thereby revealing a potential therapeutic strategy.

We then investigated whether pharmacological Polθ inhibition results in similar synergistic lethality with APOBEC3A activity. We examined the synergistic effects of ART558, a selective small-molecule inhibitor of the polymerase (catalytic) domain of Pol (60), in combination with doxycycline-inducible APOBEC3A expression using SynergyFinder (61), quantifying drug response as the cell viability percentage relative to DMSO controls (**Methods**). APOBEC3A enhanced sensitivity to ART558 with synergy scores of 10.19 in 5637 cells (p = 1.21 x 10^-2^) and 9.63 in RT112/84 cells (p = 7.98 x 10^-6^) (**Fig. 4D and 4E**). To confirm this relationship across multiple cell lines and with a different Polθ inhibitor, we analyzed data from the PRISM Repurposing project in DepMap (62) (**Methods**). Indeed, we found that urothelial cancer cells with higher APOBEC3-induced substitution burdens were significantly more sensitive to Polθ inhibition by Novobiocin, a first-in-class Polθ inhibitor (63) (Spearman r = 0.46, p = 0.01, n=23), as indicated by averaged sensitivity scores across two screens (**Fig. 4F and Supplementary Table S6**).

### Targeting Polθ induces synthetic lethality in APOBEC3A-expressing tumors *in vivo*

Having established that Pol θ inhibition induces synthetic lethality with APOBEC3A activity *in vitro*, we sought to validate this phenomenon *in vivo*. The 5637 cells (**Supplementary Fig. S3B**) with doxycycline-inducible APOBEC3A and either shPolθ or scramble shRNA were injected subcutaneously into both flanks of mice. Intraperitoneal injection of doxycycline or vehicle (PBS) was initiated when tumors reached an average of 30 mm^3^ (Day 35 post-injection) and continued every other day. Mice were randomized into four groups: scramble, shPolθ, scramble shRNA + APOBEC3A^OE^ and shPolθ + APOBEC3A^OE^. At the experimental endpoint (Day 17 post-doxycycline treatment), tumors in the shPolθ + APOBEC3A^OE^ group showed a 57% reduction in subcutaneous tumor volume compared to the scramble + APOBEC3A^OE^ group (p = 0.033). No significant difference in growth was observed between the PBS-treated scramble and shPolθ groups, indicating the specificity of the synthetic lethal interaction (**Fig. 4G**). Resected tumors showed significantly smaller mean volumes in the shPolθ + APOBEC3A^OE^ group (12.5 mm^3^) compared to scramble shRNA + APOBEC3A^OE^ controls (26.9 mm^3^) (p = 0.042) (**Fig. 4H; Supplementary Fig. S4**). Taken together, these data support the clinical potential of Polθ inhibitors in cancers with APOBEC3 activity.

## Discussion

APOBEC3-induced mutagenesis is well known to cause genomic instability and intratumoral heterogeneity, metastasis, the evolution of treatment resistance and worse clinical outcomes (15,19,64). However, no APOBEC3 inhibitors have been brought to clinical use, leaving a critical therapeutic gap (20). There is an unmet need to develop effective therapeutic strategies that leverage vulnerabilities arising from APOBEC3-induced DNA damage. Here, we show that APOBEC3A generates DSBs and then shifts the DSB repair pathway choice towards the TMEJ pathway. This occurs through the competition between APOBEC3A and RPA for ssDNA overhangs, thereby limiting RPA loading and exposing microhomology sequences that anneal to initiate TMEJ. This creates therapeutic vulnerability by depending on Polθ, the key TMEJ polymerase, thereby establishing Polθ inhibition as a rational synthetic lethal strategy for APOBEC3A-active cancers.

APOBEC3A initiates a DNA damage cascade due to the accumulation of genomic uracils resulting from cytosine deamination, which can lead to replication-fork stalling and DSB formation (25,32). The final outcomes depend on the strand and location of uracil incorporation, as well as on downstream processing mechanisms (35). APOBEC3A deaminates cytosine on both leading and lagging strands during DNA replication (34). Uracil removal by base excision repair (BER) can stall polymerase on the leading strand, causing replication stress (32) or directly generate two-ended DSBs on the lagging strand (**Fig. 4I**) (35,36). Consistent with previous findings (23,25,65-67), our data show that APOBEC3A induces CHK1 activation and replication-fork stalling, the hallmarks of replication stress. Immunofluorescence microscopy analysis showed that APOBEC3A exposure significantly increased both ssDNA and DSBs and induces S-phase arrest (45,68). This further supports a model of uracil lesions leading to replication stress, fork arrest, and, eventually, DSB formation. Mechanistically, our findings are consistent with prior observations showing that APOBEC3A competes with RPA for binding ssDNA overhangs (69) and elucidate how this competition creates conditions favoring TMEJ repair. Notably, the recruitment of Polθ by this mechanism potentially establishes a positive feedback loop in which Polθ can compete with RPA for ssDNA through its helicase domain (70), progressively shifting the repair balance toward TMEJ. Our findings provide mechanistic insight into prior observations of synthetic lethality from ATR inhibition in APOBEC3-high tumors (23,66,67,71). While ATR inhibitors exploit APOBEC3-induced replication stress, cells counter this vulnerability by activating TMEJ as the predominant repair mechanism for APOBEC3-induced DSBs. This provides a rationale for combining ATR and Polθ inhibition to prevent resistance evolution in APOBEC3-active tumors.

Our mechanistic model is supported by genomic evidence across multiple datasets. Consistent with the known role of inappropriate end-joining in generating chromosomal rearrangements such as small insertions or deletions and large-scale structural variants, driving chromosomal instability (51), our results show a significant correlation between APOBEC3-induced substitutions and higher burden of MMDs in both whole-genome data and TCGA whole-exome datasets. Independent support for this correlation comes from analysis of whole-genome sequencing data in the Pan-Cancer Analysis of Whole Genomes Project (PCAWG) (26). Furthermore, our analysis of chromosomal instability signatures revealed a strong correlation between APOBEC3A-induced substitutions and signatures of increased TMEJ activity, as evident by the microhomologies near breakpoints (52). We propose that the cascade of APOBEC3A induction followed by preferential TMEJ repair, plays a key role in driving specific patterns of chromosomal instability and potentially even in the biogenesis of specific complex structural variants including extrachromosomal DNA. Aberrant DNA repair pathways, including classical and alternative NHEJ, can produce ecDNA from chromosomal fragments (72-74). A study of urothelial bladder patient samples revealed significant upregulation of DNA repair genes, including Polθ, in ecDNA-positive tumors compared to ecDNA-negative ones(75). We recently demonstrated overlap between APOBEC3-induced mutagenesis and ecDNA biogenesis in urothelial cancer, both topographically and chronologically (19). The preferential repair of APOBEC3A-induced DSBs by TMEJ, shown here, suggests a novel role for this APOBEC3A-TMEJ axis in ecDNA formation.

While our data demonstrate APOBEC3A’s central role in promoting TMEJ, additional mechanisms may modulate this pathway. HMCES binding to abasic sites could compete with TMEJ by promoting canonical repair pathways (76,77), while persistent stress might activate complementary mechanisms that involve Polθ, such as microhomology-mediated break-induced replication (MMBIR) (78,79). While we focused our study on APOBEC3A, it is possible that other APOBEC3 family members may also influence the DSB repair pathway choice in a similar manner. Although APOBEC3A expression is generally low in most tissues, mutational signature analyses clearly demonstrate APOBEC3A-induced mutagenesis patterns across multiple tumor types, including bladder cancer (16), likely due to exposure to bursts of episodic expression during the natural history of the tumor (28,80). While we used doxycycline-inducible systems for mechanistic studies, our orthogonal data from cell lines with varying endogenous APOBEC3A levels, TCGA analysis, and DepMap datasets validate these findings in cancers with endogenous APOBEC3A activity, supporting the clinical relevance of the APOBEC3A-TMEJ interaction.

In summary, our findings position Polθ inhibition as a precision therapeutic strategy for APOBEC3-high tumors. We demonstrate synthetic lethality via both genetic knockdown and small-molecule inhibition across multiple cell line models with endogenous APOBEC3 activity, and *in vivo* validation confirms Polθ dependency under APOBEC3A-induced genotoxic stress. Mechanistically, we reveal that APOBEC3A not only generates DNA damage but actively promotes TMEJ repair through RPA displacement, creating a critical survival dependency that can be therapeutically exploited. By turning off this repair axis, Polθ inhibitors tip the balance from damage tolerance to cell death. Targeting Polθ-dependent repair in APOBEC3-active tumors, can selectively eliminate cancer cells with mutagenic APOBEC3 activity (**Fig. 4I**), thereby depleting the evolutionary reservoir of therapy-resistant variants and reducing overall tumor population fitness. The clinical development of Polθ inhibitors in Phase 1/2 trials for BRCA-deficient cancers, including ART6043, GSK4524101 and RP-3467, either alone or combined with PARP inhibitors, confirms the potential for Polθ for translation (81,82). While Polθ dependence is classically associated with HR-deficient tumors (83), our discovery that APOBEC3A-high tumors rely on Polθ via an APOBEC3A-RPA competition mechanism reveals a distinct vulnerability. It suggests a promising biomarker-based approach to extend Polθ inhibitor therapy to patients with APOBEC3A-driven cancers.

## Methods

### Cell lines

The 5637 (HTB-9), U2OS (HTB-96), and HEK293T (CRL-3216) cell lines were obtained from the American Type Culture Collection (ATCC). The RT112/84 cell line, a derivative of RT112, was obtained from Millipore Sigma (#85061106). The U2OS and HEK293T cells were cultured in DMEM (Gibco) supplemented with 100 U/ml Penicillin-Streptomycin (Gibco) and 10% tetracycline-free FBS (Omega Scientific). The 5637 cells were cultured in RPMI-1640 (Gibco) supplemented with 100 U/ml Penicillin-Streptomycin (Gibco) and 10% tetracycline-free FBS (Omega Scientific). The RT112/84 cells were cultured in EMEM (ATCC) supplemented with 100 U/ml Penicillin-Streptomycin (Gibco) and 10% tetracycline-free FBS (Omega Scientific). All the cells were maintained at 37°C in a humidified atmosphere containing 5% CO_2_. Cell line authentication was confirmed by STR profiling within the past 6 months. Cells were tested for Mycoplasma contamination using the Mycoplasma PCR Detection Kit (ABM) according to the manufacturer’s instructions. The mutational calls in 5637 and RT112 cell lines were obtained from a previous publication (29).

### Plasmids for doxycycline-inducible APOBEC3A expression

The pSLIK-APOBEC3A plasmid, a gift from Dr. Matthew D. Weitzman (25), was used to generate doxycycline-inducible APOBEC3A expression in 5637 cells. In this pSLIK-APOBEC3A-HA construct, the Venus fluorescent protein is expressed under the constitutive CMV promoter, and HA-tagged APOBEC3A is under the doxycycline-inducible Tet-On promoter. The pLVX-APOBEC3A and pLVX-APOBEC3A E72Q plasmids from Dr. Linda Chelico express FLAG-tagged APOBEC3A and its deaminase-inactive variant E72Q, respectively, under the doxycycline-inducible Tet-On promoter. These plasmids carry puromycin resistance as a selection marker and were used to generate doxycycline-inducible deaminase-active and deaminase-inactive APOBEC3A expression in RT112 cells. For the DSB reporter assay in U2OS cells, we replaced the puromycin resistance gene in the pLVX-APOBEC3A plasmid with tdTomato to enhance selection efficiency through sorting. We thereby amplified a tdTomato sequence from the pLVX-tdTomato plasmid (Takara Bio # 631238), a gift from Dr. Mohammad Kamran, using PCR. The PCR amplicon was then inserted into the pLVX-APOBEC3A backbone using T4 DNA ligase (NEB) after digestion with restriction enzymes AvrII and MluI (NEB). The resulting plasmid was validated by Sanger sequencing. Plasmids were transformed into One-Shot Stbl3 chemically competent *E. coli* cells (Thermo Fisher Scientific) according to the manufacturer’s instructions.

### Lentivirus production

Plasmids were co-transfected with packaging plasmids psPAX2 (Addgene #12259) and pCMV-VSV-G (Addgene # 8454) into HEK293T cells using Lipofectamine 2000 (Thermo Fisher Scientific). The culture supernatant containing lentiviral particles was collected every 24 hours for three days post-transfection. Lentivirus particles were filtered through a 0.45 μm filter (Corning), concentrated with Lenti-X Concentrator (TaKaRa Bio), and stored at −80°C.

### Generation of doxycycline-inducible APOBEC3A cells

Cells were transduced with concentrated lentiviral particles in the presence of 10 μg/mL polybrene (Millipore Sigma). 5637 cells transduced with pSLIK-APOBEC3A were sorted by fluorescence-activated cell sorting (FACS) for Venus-positive cells. A single-cell clone with doxycycline-inducible APOBEC3A expression was established through serial dilution. U2OS cells transduced with pLVX-APOBEC3A-tdTomato were sorted by FACS for tdTomato-positive cells. RT112/84 cells transduced with pLVX-APOBEC3A or pLVX-APOBEC3A E72Q were selected using 2 μg/mL puromycin for 48-72 hours. APOBEC3A expression levels were confirmed by SDS-PAGE Western blotting. Fluorescence-activated cell sorting (FACS) was performed at the WCM Flow Cytometry Core Facility by an experienced operator.

### SDS-PAGE western blotting

Whole-cell protein lysates were prepared using ice-cold RIPA buffer (Thermo Fisher Scientific) supplemented with protease and phosphatase inhibitor cocktails (Thermo Fisher Scientific). Protein concentrations were quantified by the Pierce BCA Protein Assay Kit (Thermo Fisher Scientific). Lysates (20-30 μg) were separated by SDS-PAGE (10% Bis-Tris, Invitrogen) using MOPS or MES running buffer and transferred to PVDF membranes using the iBlot Transfer System (Thermo Fisher Scientific). Membranes were blocked with 3% non-fat milk in 1X TBST. The following primary antibodies were incubated overnight at 4°C: anti-phospho-CHK1 (CST, 2348, 1:1000), anti-phospho-CHK2 (CST, 2197, 1:1000), anti-phospho-RPA (Sigma, PLA0071, 1:1000), anti-CHK1 (CST, 2360, 1:1000), anti-CHK2 (CST, 3440, 1:1000), anti-γH2AX (Abcam, ab11174, 1:1000), anti-cleaved PARP (Asp214) (CST, 9541, 1:1000), anti-HA-tag (CST, 2367, 1:1000), anti-FLAG-tag (Sigma, F1804, 1:1000), anti-GAPDH (CST, 2118, 1:1000), and anti-α-tubulin (Millipore, 05-829, 1:1000). HRP-conjugated secondary antibodies (goat anti-rabbit, 32260, and goat anti-mouse, 32230, Invitrogen, 1:1000) were incubated for 2 hours at room temperature. Blots were developed using Immobilon Forte Poly Western HRP Substrate (Millipore Sigma) and visualized with the ChemiDoc MP Imaging System (Bio-Rad). Membranes probed for phospho-CHK1 and phospho-CHK2 were stripped with Restore Western Blot Stripping Buffer (Thermo Fisher Scientific) and re-probed with antibodies against total CHK1 and CHK2.

### Immunofluorescence assay *in vitro* and quantification of γH2AX and RPA foci

Cells were seeded in chamber slides (Thermo Fisher Scientific) and treated with DMSO or 1 μg/mL doxycycline for 24 hours. Cells were subsequently fixed with 4% paraformaldehyde at room temperature for 10 minutes, permeabilized with 0.5% Triton X-100 in PBS, and blocked with 10% normal goat serum and 2% BSA in PBS. Primary antibodies, anti-γH2AX (CST, 9718, 1:1000) and anti-RPA2 (Thermo Fisher Scientific, MA1-26418, 1:1000), were applied overnight at 4°C. Secondary antibodies, AlexaFluor 488-conjugated donkey anti-rabbit IgG (Jackson ImmunoResearch Laboratories, 711-545-152, 1:1000) and AlexaFluor 594-conjugated donkey anti-mouse IgG (Jackson ImmunoResearch Laboratories, 715-585-150, 1:1000) were incubated for 2 hours at room temperature. Nuclei were stained with DAPI during secondary antibody incubation. Slides were mounted using Vectashield Vibrance Antifade Mounting Medium (Vector Laboratories, H-1700). Images were acquired using a fluorescence microscope (Axio Imager M2) equipped with a 63x/1.4 oil objective (Carl Zeiss). Z-stacks (0.25 μm intervals) were captured and analyzed using Fiji software. Multiple random fields of view (n=20) were selected for imaging. Images were processed by Z-projection using the “Max intensity” function in Fiji, and background subtraction was performed using the “rolling ball” algorithm for both fluorescence channels. Foci in each channel were automatically quantified using Foci-Analyzer (84). Nuclei were segmented from the DAPI channel using the StarDist algorithm on 2D maximum intensity projections, excluding edge-touching or size-outlier nuclei. Foci were detected in the γH2AX and RPA channels using a marker-controlled watershed algorithm with automated background-based thresholding and size filtering. Negative controls, omitting primary antibodies, were employed to optimize detection parameters and ensure accurate quantification. Colocalization was automatically determined by the software based on the spatial overlap of foci across channels, with nuclei containing at least one overlapping focus classified as colocalization-positive. For downstream analyses, both the total number of γH2AX foci per nucleus and the subset overlapping with RPA were recorded.

### DNA fiber assay

The DNA fiber assay was performed as previously described (85). Cells were seeded in six-well plates and treated with doxycycline (Millipore, 10592-13-9) to induce APOBEC3A expression, or with DMSO as a control, for 24 hours in serum-free medium to synchronize the cell cycle and minimize APOBEC3A-induced damage during the cell cycle (24). After 24 hours, cells were released into the cell cycle using medium containing 10% FBS. They were labeled with 25 μM CldU (MP Biomedicals, ICN10547883) for 0.5 hour, washed three times with fresh medium, and subsequently labeled with 250 μM IdU (MP Biomedicals, ICN10035701) in the presence or absence of Aphidicolin (Sigma Aldrich) at a low concentration of 0.8 μM for 3 hours to induce limited inhibition of DNA polymerase. A longer second pulse of labeling was given to allow sufficient time for uridine processing and subsequent induction of DSB repair by APOBEC3A. Labeled cells were dissociated using trypsin and harvested in cold PBS. 2 μl of cell suspension was spotted onto a glass slide and air-dried for 2 minutes, then lysed with 7 μL of fiber lysis buffer (50 mM EDTA and 0.5% SDS in 200 mM Tris-HCl, pH 7.5) for 2 minutes at room temperature. Slides were tilted at approximately 15° to allow DNA fibers to spread. Once the fiber solution reached the bottom of the slide, it was laid horizontally and air-dried. For immunofluorescence staining, slides were immersed in a methanol/acetic acid (3:1) solution for 15 minutes at −20°C, denatured in 2.5 M HCl for 30 minutes at room temperature, blocked with 5% BSA in PBS for 20 minutes, and incubated with anti-BrdU antibodies specific for CldU (Abcam, ab6326, 1:15) and IdU (BD Biosciences, 347580, 1:15) for 1 hour in a humidified chamber at room temperature. After washing three times with PBS, slides were incubated with AlexaFluor 488-conjugated goat anti-rat IgG (Invitrogen, A-11006, 1:15) and AlexaFluor 568-conjugated goat anti-mouse IgG (Invitrogen, A-11031, 1:15) antibodies for 1 hour at room temperature. Slides were washed three times with PBS and mounted with ProLong Antifade Mountant (Thermo Fisher Scientific, P36930). Images were acquired using a fluorescence microscope (Axio Imager M2) with a 63x/1.4 oil objective (Carl Zeiss). Random fields were selected for imaging, and fiber lengths were measured using ImageJ software.

### Flow cytometry for cell cycle analysis and γH2AX staining

Cells were seeded in six-well plates and treated with doxycycline to induce APOBEC3A expression or with DMSO as a control for 24 hours in serum-free medium to synchronize the cell cycle and minimize APOBEC3A-induced damage during the cell cycle (24). After 24 hours, cells were released into the cell cycle using medium containing 10% FBS and were harvested at different time points (0, 12, and 24 hours). Cells were fixed with BD Cytofix/Cytoperm solution for 30 minutes at room temperature, washed with BD Perm/Wash buffer, and stored at −4°C for subsequent staining. Before staining, the fixed cells were re-fixed using BD Cytofix/Cytoperm solution for 5 minutes at room temperature. The following primary antibodies were used for staining: Alexa Fluor 647 mouse anti-H2AX (pS139) (BD Biosciences, 5 μL per sample) and PE-conjugated anti-PCNA antibody, anti-human/mouse/rat REAfinity (Miltenyi Biotec, 2 μL per sample). Antibody incubation was performed for 1 hour at room temperature. After washing with BD Perm/Wash buffer, cells were resuspended in DAPI solution (1 μg/mL) and analyzed by flow cytometry. Data analysis was conducted using FlowJo software. Cell cycle distribution was determined using the Dean-Jett-Fox model, with a G1/G2 ratio of approximately 1.8. The percentage of cells in each cell cycle stage was calculated based on gates defined in the FlowJo workspace. Additionally, the γH2AX-positive cells in each stage were quantified.

### DNA extraction, library preparation, and whole-exome sequencing

5637 cells transduced with the pSLIK-A3A-HA vector were exposed to 2 µg/mL of doxycycline or DMSO for three days, followed by culturing in regular media for four days. This cycle was repeated six times to provide six episodic exposures of A3A activity. Genomic DNA was extracted using the DNeasy Blood & Tissue kit (Qiagen) according to the manufacturer’s instructions and stored at −80°C. DNA quality was assessed using the Agilent TapeStation system. Whole-exome sequencing libraries were prepared using the xGen Exome Research Panel v2 (IDT) at the WCM Genomics Core Facility. Pooled samples were sequenced on a NovaSeq 6000 (Illumina) system using paired-end 100bp reads (2X100 bp).

### Alignment and somatic variant calling

Short-read alignment was performed against the human reference genome (GRCh37/b37) using an in-house pipeline (10). Reads were trimmed for adapter sequences, aligned by BWA-MEM, indel realignment was performed via GATK, and PCR duplicates were removed using Picard Tools. Somatic variants were called in single-sample mode by MuTect2, Pindel, VarScan, and Vardict with default parameters. For Mutect2, variant calls were filtered in two steps. First, filtering was based on the significance scoring of each call and the estimation of sample contamination, using the CalculateContamination and FilterMutectCalls tools in GATK4. The second step involved quantifying nucleotide substitution errors caused by mismatched base pairings during various sample/library preparation stages, which were then filtered from the original MuTect2 calls using the CollectSequencingArtifactMetrics and FilterByOrientationBias tools in GATK4. Additional filtering criteria were applied as follows: 1) variants with total reads >10 and alternative allele counts >2 were retained; 2) variants called by at least two of the variant callers were considered; 3) variants present in the human dbSNP database (GRCh37p13_b151_common) were excluded using SnpEff. 4) Variants shared by both the doxycycline-treated and control groups were excluded to remove pre-existing variants in the cell line. The results from different mutation callers were combined using Python pandas based on matching genomic positions and alleles. Small insertions and deletions (indels) were retained for further analysis to identify mutational footprints generated by the DSB repair process

### Mutational footprints analysis of human tumors

Somatic variant calls and copy number segmentations of whole-genome sequencing (tumor sample n=71) were obtained from our published study as previously described (19). The HR deficiency score was calculated using HRDetect on the whole-genome sequencing data (86). Four samples with an HRDetect probability over 70% were excluded to prevent confounding effects due to increased reliance on TMEJ in HR-deficient samples (**Supplementary Table S7**). Whole-exome data from TCGA pan-cancer cohort (n=8836) were sourced from cBioPortal. Single-base substitution and indel matrices were generated using SigProfilerMatrixGenerator (47). APOBEC-induced mutations were defined as C to T/G substitutions in the TCW (W=A/T) motifs. Deletions containing microhomology sequences were considered as indel footprints potentially associated with TMEJ.

### Distance between mutational clusters and indels

Substitution clusters were identified using SigProfilerClusters as previously described (19,87). Each substitution cluster was categorized based on its mutations. An APOBEC3-induced substitution cluster was defined when coordinated C to T/G substitutions in the TCW (W=A/T) motifs constituted greater than or equal to two-thirds of the total substitutions within the cluster. Clusters that did not meet this criterion were considered induced by other mutagenic factors. To investigate the spatial relationship between substitution clusters and indel footprints, we combined the genomic coordinates of substitution clusters and indel footprints by selecting those where the distance between the cluster center and the indel footprint was less than 0.1Mb using a custom Python script. We then calculated the genomic distances between substitution clusters and their corresponding indel footprints. Mean distances were calculated separately for APOBEC3-induced substitution clusters and clusters induced by other mutagenic factors. These distances were analyzed to assess whether indel footprints are closer to APOBEC3-induced clusters than to other clusters.

### Chromosomal instability signatures

Chromosomal instability signatures for our 67 tumor samples were quantified using the CINSignatureQuantification package (52). Chromosomal instability signature matrices for TCGA pan-cancer cohorts were obtained from the data repository of the Drews et al. paper (52) (https://github.com/markowetzlab/Drews2022_CIN_Compendium).

### DSB repair reporters

The pDR-GFP reporter was a gift from Maria Jasin (Addgene, #26475). The EJ2GFP-puro reporter was a gift from Jeremy Stark (Addgene, #44025). These plasmids were transfected into flow-sorted U2OS cells expressing pLVX-APOBEC3A-tdTomato using Lipofectamine 3000 (Thermo Fisher Scientific) following the manufacturer’s instructions. Transfected cells were selected with puromycin (2 μg/mL) for more than 2 weeks before the experiments. Selected cells were seeded in six-well plates and treated with 1 μg/mL doxycycline to induce APOBEC3A expression or with DMSO as a control for 24 hours in serum-free medium to synchronize the cell cycle and minimize APOBEC3A-induced damage during the cell cycle (24). After 24 hours, adeno-associated virus (AAV) particles containing I-SceI were added to the medium containing 10% FBS to induce DSBs. After 36 hours, the cells were harvested for flow cytometry analysis to detect GFP-positive cells.

### Amplification of adeno-associated virus (AAV) encoding I-SceI

Dr. Kent Mouw provided frozen AAV particles encoding the I-SceI nuclease. The AAV particles were added to a T75 flask of HEK293T cells. The cells were incubated for 3 days post-transduction and then harvested. After washing the cell pellet with PBS, resuspend the cells in 500 μL of 10 mM Tris-HCl, pH 8.0. AAV particles were released from cells by performing three rapid freeze-thaw cycles using a 100% ethanol-dry ice bath (freezing) and a 37°C water bath (thawing). Cellular debris was removed by centrifuging at 1500 rpm for 10 minutes at 4°C. The supernatant containing AAV particles was stored at −80°C in aliquots.

### Protein expression and purification for biochemical assays

Recombinant baculovirus production of GST-tagged APOBEC3A was performed using the BD Biosciences pAcG2T transfer vector system as previously described (88,89). Purification of GST-APOBEC3A and cleavage from the GST tag was performed as previously described (90). The bacterial expression plasmid p11d-tRPA, containing the three subunits of human RPA, was a gift from Dr. Marc Wold. Expression and purification of RPA from *E. coli* were performed as previously described (91). The Sumo3-PolQM1 plasmid (Addgene, #78462), a gift from Drs. Sylvie Doublie and Susan Wallace, was used to express the polymerase domain of Polθ (Polθ-pol) (58). For purification of Polθ-pol, 8 liters of Rosetta2(DE3) pLysS cells containing the Sumo3-PolQM1 plasmid were grown in Terrific Broth without α-lactose and induced with 1mM IPTG for 16 hours at 23°C. Cells were recovered from the broth, resuspended in lysis buffer (50 mM HEPES, pH 8.0, 300 mM NaCl, 10% glycerol, 20 mM imidazole, and 5 mM CaCl_2_), frozen in liquid nitrogen, and stored at −80°C until use. Before cell disruption, 200 mL of total lysis solution (1.5% NP-40 Alternative, 5 mM β-mercaptoethanol, EDTA-free protease inhibitor cocktail, 10 mM PMSF, 100 mM benzamidine, 5 mM MgCl_2_, 100 mg lysozyme, 500 U DNaseI, and 5 KU Benzonase) was added to the defrosted cell suspension. Cells were disrupted using a cell disruptor operated at 35,000 psi, and then the lysate was cleared twice by centrifuging at 27,000 x g for 30 min at 4°C. The purification was then performed using ice-cold buffers. First, the cleared lysate was applied to a 5 mL His-Trap column in buffer (50 mM HEPES, pH 8.0, 300 mM NaCl, 10% glycerol, 5 mM β-mercaptoethanol, 0.005% NP-40, and 20 mM imidazole). The column was washed with the loading buffer and then eluted with the loading buffer containing 500 mM imidazole. The Polθ-pol containing fractions were then applied to a ceramic hydroxyapatite column and subsequently to a heparin column, as previously described (58). Following heparin column purification, the Polθ-pol fractions were desalted using a Bio-Gel P-6 10mL desalting column with buffer (50 mM HEPES, pH 7.0, 10 mM NaCl, 10% glycerol, 5 mM β-mercaptoethanol, and 0.005% NP-40). Then they were loaded onto a HiTrap SP column equilibrated with low-salt buffer (50 mM HEPES, pH 7.0, 10 mM NaCl, 10% glycerol, 5 mM BME, and 0.005% NP-40 substitute). Elution was performed using a salt gradient, with Polθ-pol eluting at 450 mM NaCl. Polθ-pol was then applied to a Superose 12 column equilibrated with buffer containing 50 mM HEPES, pH 8.0, 300 mM NaCl, 10% glycerol, 5 mM BME, and 0.005% NP-40 substitute. Fractions containing the ∼100 kDa protein were collected. Individual fractions were tested for nuclease activity, and only those lacking nuclease activity were used in the biochemical assay for TMEJ.

### Biochemical assay for TMEJ

A DNA substrate (F1/S2, **Supplementary Table S8**) was formed by annealing two DNA complementary oligonucleotides at a 1:1.2 ratio in 50 mM Tris (pH 7.5) and 50 mM NaCl. The F1 oligonucleotide was labeled at the 5’ end with fluorescein. The reaction was carried out in TMEJ buffer containing 25mM Tris-HCl (pH 8.8), 1mM DTT, 0.01% NP-40 Alternative, 0.1mg/mL BSA, 3% glycerol, 10mM MgCl_2_, 2mM ATP, 20µM dNTPs, and 30mM NaCl. The reaction mixture included 100nM F1/S2 DNA substrate, 50nM Polθ-pol, 50nM RPA, and 100nM APOBEC3A. Glycerol was added to a final concentration of 4% to facilitate gel loading. The reaction was initiated by adding the F1/S2 DNA substrate and incubated at 37°C. Samples were taken at 2, 5, and 10 minutes, then mixed with an equal volume of stop buffer containing 100 mM Tris-HCl (pH 7.5), 10mg/mL proteinase K, 80 mM EDTA, and 0.5% SDS, followed by incubation at 37°C for 20 minutes. DNA samples were directly loaded onto an 11% non-denaturing polyacrylamide gel without adding loading dye and resolved at 4°C. Gels were scanned using a Chemidoc MP Imaging System (Bio-Rad) and analyzed using ImageQuant software (GE Healthcare). ImageQuant was used to measure integrated band intensities within each lane. Background subtraction was performed using the no-enzyme control lane (DNA oligonucleotides only), and background was subtracted from a position equivalent to the product band. The intensities were normalized so that the total intensity in one lane equaled 100%, and the values were distributed accordingly between the substrate and product bands.

### shRNA knockdown of Polθ

The shRNA plasmid targeting Polθ (pLV-Bsd-U6-Polθ shRNA) and a scrambled shRNA control (pLV-Bsd-U6-scramble shRNA) were generated by VectorBuilder. The target sequence for Polθ knockdown is listed in **Supplementary Table S8**. Plasmids were validated by Sanger sequencing. Knockdown efficiency was confirmed by quantitative real-time PCR.

### RNA extraction and quantitative PCR

Total RNA was extracted using the RNeasy Plus Mini Kit protocol (Qiagen) according to the manufacturer’s instructions. RNA concentration was determined by Varioskan™ LUX multimode microplate reader (Thermo Fisher Scientific). cDNA was synthesized using the SuperScript III First-Strand Synthesis System (Invitrogen). Quantitative PCR was performed using the QuantStudio real-time PCR system (Thermo Fisher Scientific) with Power SYBR Green Master Mix (Applied Biosystems). All reactions were run and analyzed using QuantStudio PCR software (Thermo Fisher Scientific). Expression levels of Polθ were normalized to *GAPDH*. Primers sequences are listed in **Supplementary Table S8**.

### Cell viability and synergy score

Cell viability was measured using 384-well microplates (Corning). The CyQUANT™ Direct Cell Proliferation Assay (Thermo Fisher Scientific) was used to quantify cell viability, following the manufacturer’s instructions. Measurements were taken with a Varioskan™ LUX multimode microplate reader (Thermo Fisher Scientific). For viability assay of cells with Polθ knockdown, 2,000 cells were seeded into wells, and after 24 hours, either DMSO or doxycycline was added. After 2 days of treatment, cell viability was measured. For experiments with RT112/84 cells to calculate synergy scores, varying concentrations of Polθ inhibitor and doxycycline were mixed and added to the cells. Each condition had at least six replicates. After 4 days of treatment, cell viability was measured. Viability was normalized to the DMSO controls. For experiments with 5637 cells to calculate synergy scores, the cells were seeded into a 96-well plate.

Varying concentrations of Polθ inhibitor and doxycycline were mixed and added to the cells. Then, the cells were traced by live-cell imaging using the Incucyte system (Sartorius) for 4 days. Each condition has two independent wells, and each well has four imaging as replicates. Cell confluences were calculated using Incucyte software. Synergistic score analysis was performed using SynergyFinder Plus (61), based on the confluence matrix. The synergistic score was determined using the ZIP model using optimized continuous diluted concentrations of doxycycline and inhibitors. The p-value was calculated by comparing the observed synergy scores with a null distribution generated from a non-interaction model, allowing us to assess the statistical significance of the synergistic effects using SynergyFinder Plus (61).

### Genetic dependency score and Novobiocin sensitivity data from CCLE

Genetic dependency scores were obtained from the DepMap project (Chronos gene dependency probability, 24Q2 release) (59,92). Novobiocin sensitivity data were obtained from the DepMap project (PRISM Repurposing Public 24Q2). The average sensitivity value from two rounds of screening (Primary and secondary screens) was used for the final analysis (Novobiocin 2.5 µM, BRD-K85307935-236-09-5). Mutation data for urothelial carcinoma cell lines were obtained from cBioPortal. Single-base substitution matrices were generated using SigProfilerMatrixGenerator (47). APOBEC3-induced substitutions were defined as C to T/G substitutions in the TCW (W=A/T) motifs.

### Xenograft experiments

Thirty 4-week-old male NOD.Cg-Prkdcscid Il2rgtm1Wjl/SzJ (NSG) mice (Jackson Laboratory), maintained in-house by the WCM mouse facility, were used for *in vivo* experiments. Mice were housed in pathogen-free conditions with access to food and water.

Early passage 5637 cells containing doxycycline-inducible APOBEC3A were confirmed mycoplasma-negative (Lonza PCR kit) before injection. Mice were anesthetized with isoflurane and injected subcutaneously in both flanks with 1 × 10^6^ cells in 100μl PBS, using a 27G needle. Mice were randomized in two steps. Before cell injection, mice were assigned to the scramble (n=16) or shPolθ (n=14) groups, stratified by body weight and cage location. After tumors became detectable, each group was further subdivided into doxycycline (APOBEC3A overexpression) and vehicle (PBS) treatment cohorts, ensuring comparable baseline tumor volume (∼30 mm^3^) before APOBEC3A induction. Final groups assigned were scramble (PBS, n=8), shPolθ (PBS, n=7), scramble+APOBEC3A^OE^ (Doxycycline with APOBEC3A overexpression, n=8), and shPolθ+APOBEC3A^OE^ (Doxycycline with APOBEC3A overexpression, n=7). Power analysis (G*Power 3.1) indicated that n=7–8 mice per group provided 80% power (α=0.05) to detect significant differences. Doxycycline hyclate powder (Sigma-Aldrich) was dissolved in 1x PBS and administered intraperitoneally at 50 mg/kg (93) in 100 μL following a 2-day, 2-day, and 3-day interval regimen, starting on day 35 post-implantation, when tumors reached ∼30 mm^3^. Injections were administered between 1 PM and 3 PM, alternating abdominal quadrants. Control groups received PBS vehicle only. Tumors were measured weekly with a digital precision caliper once bilateral tumors were visible. Length (longest dimension) and width (shortest dimension) were measured under isoflurane anesthesia. Measurements were performed under blinded conditions. Investigator 1 (blinded) measured tumor dimensions, while investigator 2 handled the mice and recorded the corresponding animal IDs. Tumor volume “mm^3^” was calculated using the modified ellipsoid formula: (minimum diameter)^2^ × maximum diameter / 2 (94). Tumors <1 mm in any dimension were recorded as not measurable (NA). Mice were sacrificed at day 56 (Day 17 post-doxycycline treatment) when control tumors reached around 150 mm^3^. The study was terminated before reaching the IACUC-approved maximum allowable tumor. Termination criteria were significant tumor regression in the doxycycline-treated groups, and mean weight loss approaching the IACUC-mandated limit. Euthanasia was performed using CO_2_ inhalation, followed by cervical dislocation in compliance with IACUC guidelines.

## Data and Code Availability

The whole-exome sequencing files for the 5637 cell line experiments are deposited in the Sequence Read Archive with the accession number SUB13856242 (PRJNA1184822) (https://dataview.ncbi.nlm.nih.gov/object/PRJNA1184822?reviewer=c6sqtjbeap4j3lp58894hthcok). This dataset has also been analyzed for different purposes in another study by our group focusing on single-base substitutions and intratumor diversity. The current manuscript examines the indel footprints and copy number alterations induced by APOBEC3A. Resource data illustrating raw, uncropped western blots, immunofluorescence images for RPA and γH2AX, DNA fiber assay imaging data, custom script used to combine the substitutional clusters and indel footprints based on a certain distance, and HRDetect WGS data have been deposited in Zenodo (10.5281/zenodo. 13988922). The TCGA pan-cancer human cancer data and pan-cancer cell line data were obtained from cBioPortal. The Chronos gene dependency score and PRISM Repurposing dataset for Novobiocin sensitivity were obtained from DepMap (24Q2). The chromosomal instability signatures data for TCGA pan-cancer cohorts were obtained from Signature_Compendium_v5_Cosine_0.74_Acti vities_raw_THRESH95_NAMESAPRIL21.rds in the repository for Drew et al.’s paper (52). The open-source code used in this paper includes: ComplexHeatmap (https://bioconductor.org/packages/release/bioc/html/ComplexHeatmap.html), BWA MEM (https://github.com/lh3/bwa), GATK (https://github.com/broadinstitute/gatk), Picard (https://github.com/broadinstitute/picard), MuTect2 (https://github.com/broadinstitute/gatk), Pindel (https://github.com/genome/pindel), VarScan (https://github.com/dkoboldt/varscan), VarDict (https://github.com/AstraZeneca-NGS/VarDict), SnpEff (http://pcingola.github.io/SnpEff/), SigProfilerMatrixGenerator (https://github.com/AlexandrovLab/SigProfilerMatrixGenerator), SigProfilerClusters (https://github.com/AlexandrovLab/SigProfilerClusters), HRDetect (https://github.com/Nik-Zainal-Group/signature.tools.lib), SynergyFinder plus (https://synergyfinder.org), CINSignatureQuantification (https://github.com/markowetzlab/CINSignatureQuantification), Foci-analyzer (https://github.com/BioImaging-NKI/Foci-analyzer).

## Ethics Statement

All experimental procedures, animal care, and use were strictly conducted in compliance with institutional guidelines and approved by the Weill Cornell Medical College Institutional Animal Care and Use Committee (IACUC Protocol 2017-0048).

## Author Contributions

Initiation and design of the study: W.L., B.M.F. Experimental work: A.B., W.L., P.Y., A.S., M.T.B., M.D., M.O., D.N., A.B., J.G., K.W.M., M.G.R., L.C. Statistical and bioinformatics analyses: W.L., A.B., B.B., W.H., O.E., N.R, M.O. Writing of the first draft of the manuscript: W.L., A.B., and B.M.F. All authors contributed to the writing and editing of the manuscript and approved the manuscript.

## Funding

This work was supported by the National Institutes of Health NIH R37 CA279737-01A1.

A.S. was supported by an Undergraduate Student Research Award from the Natural Sciences and Engineering Research Council of Canada (NSERC) and the University of Saskatchewan, College of Medicine Biomedical Summer Research Project Award.

L.C. was supported by the Canadian Institutes of Health Research Project Grant (PJT-159560).

## Acknowledgments

We want to express our gratitude to the staff at the Weill Cornell Genomics Resources Core Facility and Flow Cytometry Core Facility. We thank Dr. Abby Green and Dr. Matthew D. Weitzman for providing the doxycycline-inducible APOBEC3A plasmid. We also thank the University of Saskatchewan College of Medicine Protein Characterization and Crystallization Facility for supplying the necessary equipment and resources for purifying Polθ-pol. ChatGPT and Claude were utilized for proofreading and editing to enhance the readability and clarity of the text.

## Competing Interests

The authors declare no competing interests.

**Supplementary Figure S1. Schematic model for the formation of DSB induced by APOBEC3A**. Breaks occur in the template strand while lagging-strand synthesis resumes in the APOBEC3A-loaded replication fork, leaving intact Okazaki fragments flanking the break, generating daughter DNA with two-ended DSBs.

**Supplementary Figure S2. APOBEC3A expression causes replication-fork stalling and cell cycle arrest in S phase**. A, Representative DNA fiber images from cells exposed to APOBEC3A or DMSO control, showing altered replication dynamics. B, Quantification of the IdU to CldU length ratio in 5637 cells illustrates a significant reduction in the length of nascent DNA fragments in APOBEC3A-expressing cells compared to DMSO control. Aphidicolin-treated cells serve as control for replication inhibition. 80-100 fibers per condition were quantified from two independent experiments. A Box plot with individual data points shows median with interquartile range, and the whiskers span from minimum to maximum value. One-way ANOVA with multiple comparisons. *: p < 0.05. C, Bar chart illustrating the distribution of cells across different cell cycle stages showing enrichment in S-phase cells induced by APOBEC3A.

**Supplementary Figure S3. APOBEC3A induction in the U2OS cell line and shRNA mediated Polθ inhibition**. A, SDS-PAGE western blotting showing the inducible expression levels of APOBEC3A-Flag in U2OS cells. B, Efficiency of Polθ knockdown (2.5 fold) using shRNA in 5637 cells with doxycycline-inducible APOBEC3A, measured by quantitative PCR. Each dot represented one replicate. Two-sided t-test. Data represent mean ± SD

**Supplementary Figure S4. Representative resected macroscopic tumor images from scramble and shPolθ xenografts with and without APOBEC3A**. Representative images of 3 resected tumors from each experimental group (scrambled, shPolθ, scramble+APOBEC3A^OE^, and shPolθ +APOBEC3A^OE^ at study endpoint (day 56). shPolθ +APOBEC3A^OE^ shows the smallest size among groups. Scale placed beside tumor during taking picture (millimeters) for reference.

**Supplementary Table S1. Indel matrices for 5637 cells intermittently exposed to DMSO and doxycycline**.

**Supplementary Table S2. Single-base substitution and indel matrices in 67 urothelial tumors based on the whole-genome sequencing data**.

**Supplementary Table S3. Combined substitution clusters and corresponding indel footprints in 67 urothelial tumors**.

**Supplementary Table S4. Counts of chromosomal instability signature contributions in 67 urothelial tumors**.

**Supplementary Table S5. Chronos Polθ dependency scores and APOBEC3-induced substitution burdens in urothelial cancer cell lines**.

**Supplementary Table S6. Sensitivity score to Novobiocin and APOBEC3-induced substitution burdens in urothelial cancer cell lines**.

**Supplementary Table S7. HR deficiency score for a total of 71 urothelial tumors**.

**Supplementary Table S8. Sequences of oligonucleotides, shRNA targeting Polθ, and primers used in this study**.

## Notes

### Competing Interest Statement

The authors have declared no competing interest.

