## Supplementary figures and images for "APOBEC3A-Induced DNA Damage Drives Polymerase θ Dependency and Synthetic Lethality in Cancer"

### Supplementary Figure S1

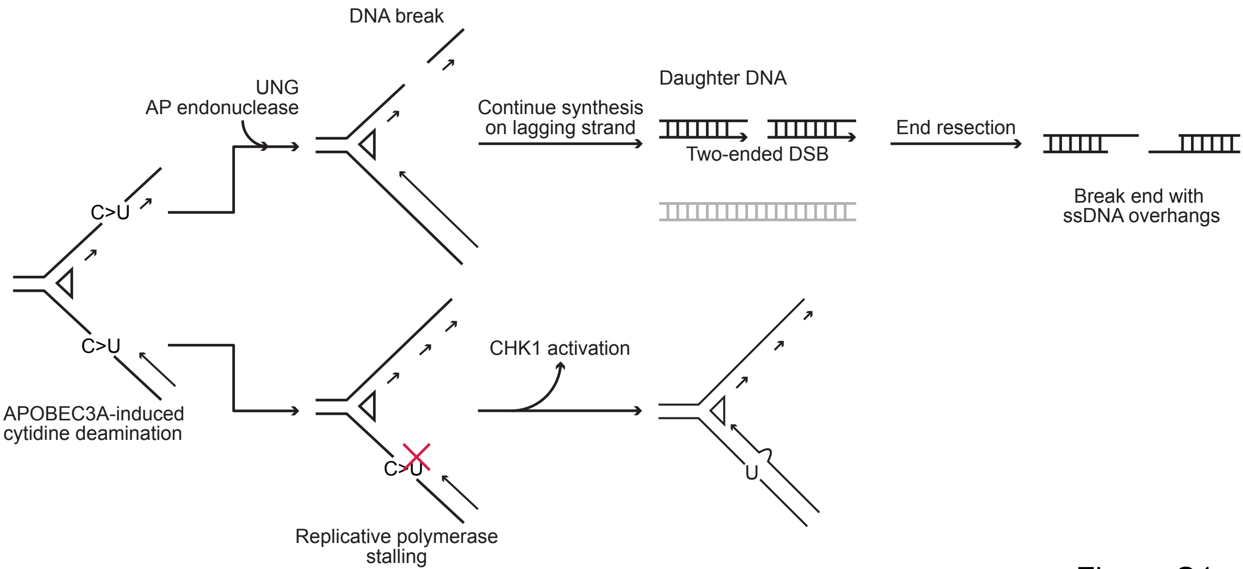

### Supplementary Figure S2

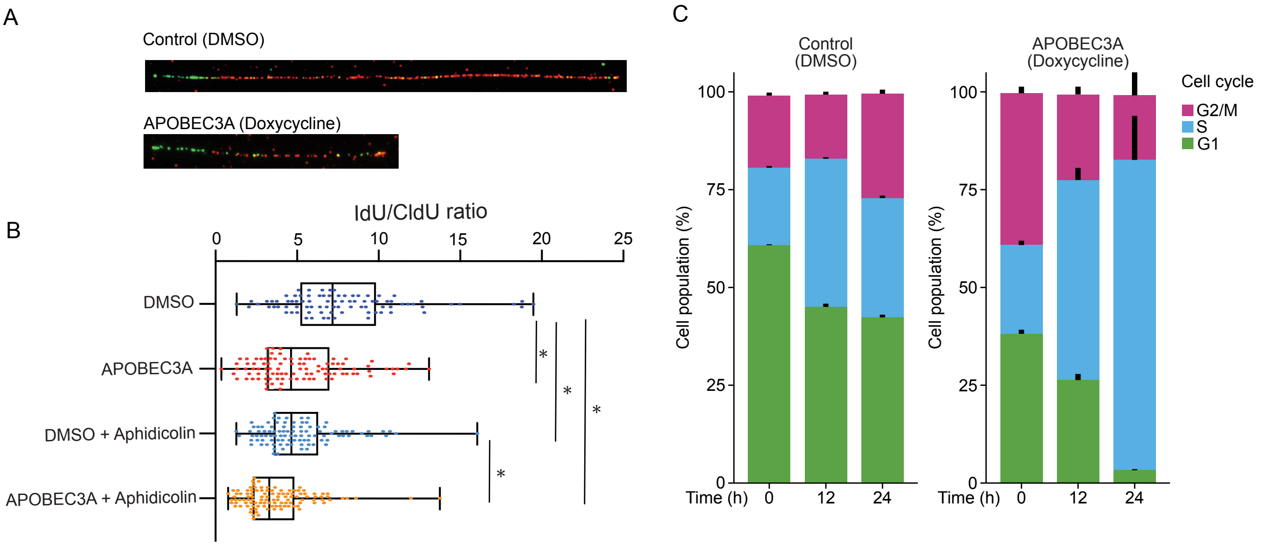

### Supplementary Figure S3

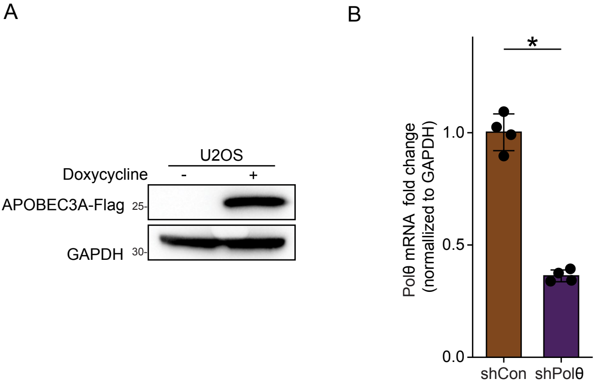

### Supplementary Figure S4

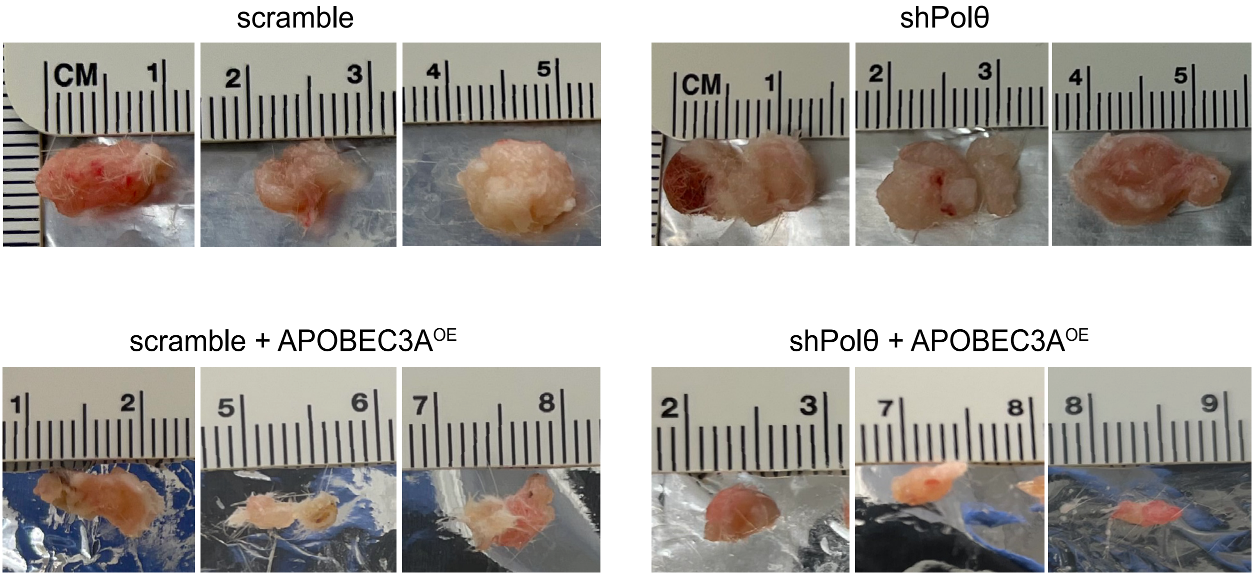


Figure S4
